# Unveiling the functional fate of duplicated genes through expression profiling and structural analysis

**DOI:** 10.1101/2024.10.29.620890

**Authors:** Alex Warwick Vesztrocy, Natasha Glover, Paul D. Thomas, Christophe Dessimoz, Irene Julca

**Affiliations:** SIB Swiss Institute of Bioinformatics, 1015 Lausanne, Switzerland; Department of Computational Biology, University of Lausanne, 1015 Lausanne, Switzerland; BioSoft Research UK, 3rd Floor, 86-90 Paul Street, London, EC2A 4NE, United Kingdom; Department of Population and Public Health Sciences, University of Southern California, Los Angeles, USA

**Keywords:** least diverged orthologue conjecture, duplication, structure, expression

## Abstract

Gene duplication is a major evolutionary source of functional innovation. Following duplication events, gene copies (paralogues) may undergo various fates, including retention with functional modifications (such as sub-functionalisation or neo-functionalisation) or loss. When paralogues are retained, this results in complex orthology relationships, including one-to-many or many-to-many. In such cases, determining which one-to-one pair is more likely to have conserved functions can be challenging. It has been proposed that, following gene duplication, the copy that diverges more slowly in sequence is more likely to maintain the ancestral function —referred to here as “the least diverged orthologue (LDO) conjecture”. This study explores this conjecture, using a novel method to identify asymmetric evolution of paralogues and apply it to all gene families across the Tree of Life in the PANTHER database. Structural data for over 1 million proteins and expression data for 16 animals and 20 plants were then used to investigate functional divergence following duplication. This analysis, the most comprehensive to date, revealed that whilst the majority of paralogues display similar rates of sequence evolution, significant differences in branch lengths following gene duplication can be correlated with functional divergence. Overall, the results support the least diverged orthologue conjecture, suggesting that the least diverged orthologue (LDO) tends to retain the ancestral function, whilst the most diverged orthologue (MDO) may acquire a new, potentially specialised, role.

## Introduction

Gene duplication is widely recognised as a key mechanism driving genome evolution (Ohno 1970). Indeed, redundant gene copies provide a *“temporary escape from the relentless pressure of natural selection”* (Ohno 1972), allowing the evolution of new functions. These genes that result from duplication events are referred to as “paralogues”, in contrast to “orthologues”, which descend from a common ancestral gene (Fitch 1970).

The role of duplication in the emergence of new gene functions has been extensively discussed (Hurles 2004). It is generally accepted that orthologues tend to be more conserved in function compared to paralogues – a concept known as the “orthologue conjecture” (Nehrt et al. 2011; Altenhoff et al. 2012; Gabaldón and Koonin 2013). Following a duplication event, the two gene copies can be retained with functional redundancy, diverge (sub- or neo-functionalization), or become lost (Force et al. 1999; Sémon and Wolfe 2007; Leitch and Leitch 2008). The most common phenomenon is that one copy accumulates deleterious mutations and thereby is silenced and eventually pseudogenised. Alternatively, mutations can lead to functional divergence between the duplicates, resulting in subfunctionalisation (each copy may retain part of the ancestral function), or neofunctionalisation (one copy may retain the ancestral function, while the other acquires a new function) (Lynch and Force 2000; Adams and Wendel 2005; Sehrish et al. 2014). Thus, gene duplication can provide the raw material for the evolution of functional innovation.

Asymmetric sequence evolution of paralogues can correlate with asymmetric functional divergence. It is generally accepted that following duplication, the two gene copies typically undergo a period of accelerated evolution (Huminiecki and Wolfe 2004), which may be apparent only in one copy (Huerta-Cepas et al. 2011; Pegueroles et al. 2013; Pich I Roselló and Kondrashov 2014). This asymmetry has been interpreted as supporting the Ohno model of evolution (Scannell and Wolfe 2008; Pegueroles et al. 2013), which hypothesises that one copy (“slow” copy, shorter branch in a gene tree) maintains the ancestral rate of evolution and function, while the other copy (“fast” copy, longer branch) may acquire a novel function (Ohno 1970).

The orthologue conjecture has motivated the notion of a “least diverged orthologue” (LDO) in the PANTHER database of phylogenetic trees (Mi et al. 2010). To define subfamilies, branch lengths are compared for each lineage following duplication, and only the subtree with the shortest branch is retained as the LDO subfamily. In this way, a single orthologous pair can be chosen even in the presence of post-speciation duplication events, with the pair being expected to be more likely than other pairs to retain the ancestral function. The advantage of this approach is that the least diverged orthologues are distinguishable in all cases, except those where the branch lengths following duplication are identical. The disadvantage, however, is that even small amounts of stochastic variation in the distance estimation will be conflated with actual differences in evolutionary rates.

Here, a generalisation of this approach is used: in instances where gene duplications arise following a speciation event, orthologous relationships can take the form of one-to-many or many-to-many. In this study, the copy with the shorter branch in the gene tree is termed the “least diverged orthologue” (LDO), while the copy with the longer branch is called the “most diverged orthologue” (MDO). Furthermore, the notion in PANTHER that LDO gene pairs are more likely to retain the ancestral function can also be applied to this framework: after duplication, the gene copy with the shorter branch (LDO) is more likely to maintain the ancestral function than that with the longer branch (MDO). Although previous work did not use this terminology, the notion that the LDO tends to retain ancestral function remains an open question. This is referred to here as the “Least Diverged Orthologue Conjecture”.

Several studies have investigated aspects of the LDO conjecture empirically. Some mainly focused on the “orthologue conjecture”, while others investigated the asymmetrical evolution of paralogues. These studies have identified a distinct rate of sequence evolution between paralogues, indicative of functional differences (Huminiecki and Wolfe 2004; Cusack and Wolfe 2007; Scannell and Wolfe 2008; Pegueroles et al. 2013; Pich I Roselló and Kondrashov 2014). To assess functional divergence between paralogous genes, various datasets have been employed, including Gene Ontology (GO) terms (Nehrt et al. 2011; Altenhoff et al. 2012), protein-protein interactions (Bandyopadhyay et al. 2006; Kim and Yi 2006), and expression patterns (Gu et al. 2005; Li et al. 2005; Wang et al. 2012; Assis and Bachtrog 2013). Additionally, the fast-evolving copy has been associated with increased levels of tissue specificity (Scannell and Wolfe 2008; Huerta-Cepas et al. 2011). Furthermore, some studies have demonstrated that even the slower evolving copy shows evidence of a burst of protein sequence evolution immediately after duplication (Scannell and Wolfe 2008), whilst others have observed an overall increase in evolutionary rate post-duplication not typically associated with rate asymmetry (Vance et al. 2022). However, the majority of these studies have relied on sequence differences (*K*_a_/*K*_s_, *d*_N_/*d*_S_) to differentiate between the fast- and slow-evolving copies, along with expression data from a restricted set of species and anatomical samples.

To investigate the LDO conjecture, this study aims to develop a novel method for capturing differences in selective pressures by identifying significant rate shifts in branch lengths following gene duplication events. Additionally, it seeks to explore functional differences between evolutionary models with statistically significant rate differences by analysing protein structural data for over 1 million proteins and integrating mRNA expression data from 16 animal and 20 plant species. Finally, the study examines the specialisation hypothesis, which proposes that the least diverged orthologue typically retains ancestral function, while the other copy evolves into a more divergent and specialised role.

## Results

### Least diverged orthologue (LDO) conjecture analysis across the Tree of Life

This study investigated the least diverged orthologue (LDO) conjecture, which implies that the gene copy with the shortest branch in the gene tree tends to retain the ancestral function after a duplication event. Analysis was performed on a dataset comprising 143 organisms representing different lineages of the Tree of Life, across 15,693 gene trees. This conjecture relies solely on observed branch length and classifies gene copies with the shorter branch as the least diverged orthologue (LDO) and those with the longer branch as the most diverged orthologue (MDO).

However, when gene families have different evolutionary rates, it is not clear whether small differences in branch length estimates can accurately classify gene copies into these two categories (LDO / MDO). Here, in order to detect distinct evolutionary rates on each of the branches, a statistical method was developed that allows for these family-specific evolutionary rates. This method is able to identify branches in a gene tree with accelerated rates of protein sequence evolution using only branch length information. This offers two key advantages: (1) it allows us to leverage precompiled libraries of gene trees such as PANTHER, eliminating the need to construct a custom dataset, and (2) it enables the analysis of ancient gene duplication events, where nucleotide substitutions have reached saturation and the usual approach of using *K*_a_/*K*_s_ cannot be reliably applied (Smith and Smith 1996; Gharib and Robinson-Rechavi 2013). Additionally, this method assigns an expected branch length to each gene tree branch, providing a model-based score that reflects deviations from the typical evolutionary rate of the gene family. As a result, this method attempts to classify a branch as typical, short, or long relative to the expected rate for the gene family.

To capture differences in evolutionary rates across branches in each gene tree, the method estimates gene family-specific rates of evolution (family-specific factors) for each gene tree (Fig. 1A). A factor greater than one indicates that a gene family is evolving more rapidly than average, potentially reflecting relaxed functional constraints, whilst a factor below one suggests slower evolution, which may reflect stronger functional constraints. The distribution of these factors has a mean of 1.06 and a standard deviation of 0.76 (Supplemental Fig. S1, Supplemental Table S1). A slight skew toward gene families with factors below one suggests that a considerable number may be evolving under stronger evolutionary constraints.

**Figure 1.**
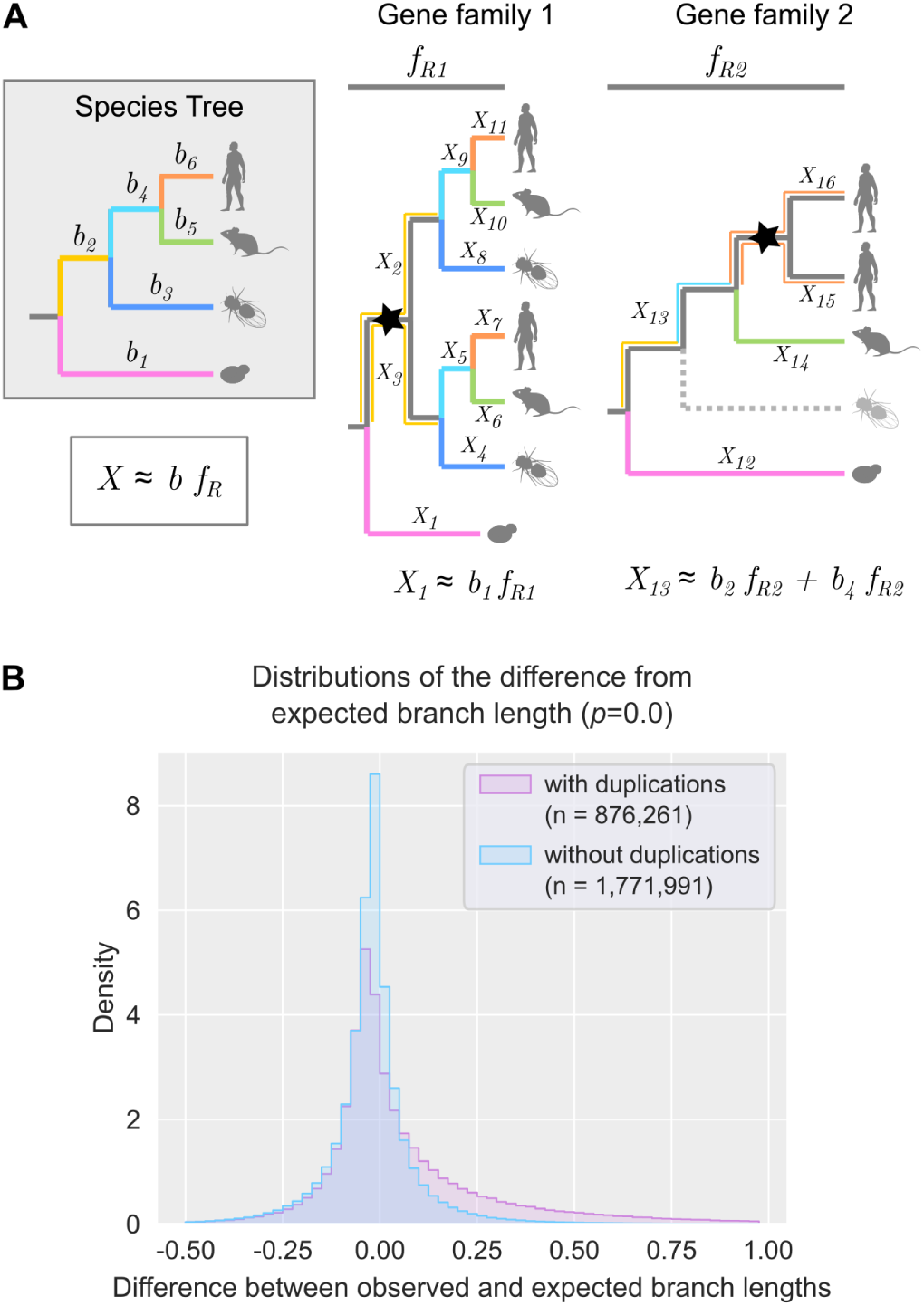
Analysis and calculation of gene tree expected branch lengths. **(A)** Graphical illustration of the calculation of the expected branch lengths (*X*). On the left, a species tree depicts the phylogenetic relationships of four species with branches highlighted by different colours and branch numbers (*b*_1_ to *b*_6_). On the right, two gene trees show a duplication event (node marked with a star). The first gene tree is complete, whilst the second tree has a gene loss represented by a dashed line. *f*_R_ denotes the family-specific rate. When no losses occur (Gene family 1), there is an exact correspondence between the speciation branch in the gene tree and the species tree (*X*_1_ to *X*_11_). When there is a loss (Gene family 2), the speciation branch in the gene tree represents the sum of the corresponding branches in the species tree (*X*_13_). Note that in both gene families, when a duplication occurs, all paralogous branches are calculated, and the expected values remain the same. **(B)** Distributions of the differences from expected branch lengths in branches with duplications (pink) and without duplications (light blue). The sample size (n) is indicated in the figure. The *p*-value for the Mann-Whitney *U* test is shown between brackets (*p*=0.0).

Using the family-specific factors, the expected branch length was computed for all branches in every gene tree (Fig. 1A) before computing the *Z*-score of the differences between the observed and expected branch lengths. A significant increase in evolutionary rate was observed between branches with no duplication (average −0.02) and those with duplications (0.07) (*p*<0.001 Mann-Whitney *U*, Fig. 1B), suggesting that paralogues tend to evolve faster than orthologues. Similar results were observed when each major lineage was analysed independently (Supplemental Fig. S2).

Following this analysis, duplication events were identified across all gene trees and classified into six categories (Fig. 2A, Supplemental Table S2): i) normal-normal: both branches have a typical rate of evolution and are consistent with their expected lengths, ii) short-short: both branches are significantly shorter than expected, iii) long-long: both branches are significantly longer than expected, iv) normal-long: only one branch is significantly longer than expected, v) short-normal: only one branch is significantly shorter than expected, vi) short-long: one branch is significantly shorter than expected and the other is significantly longer than expected.

**Figure 2.**
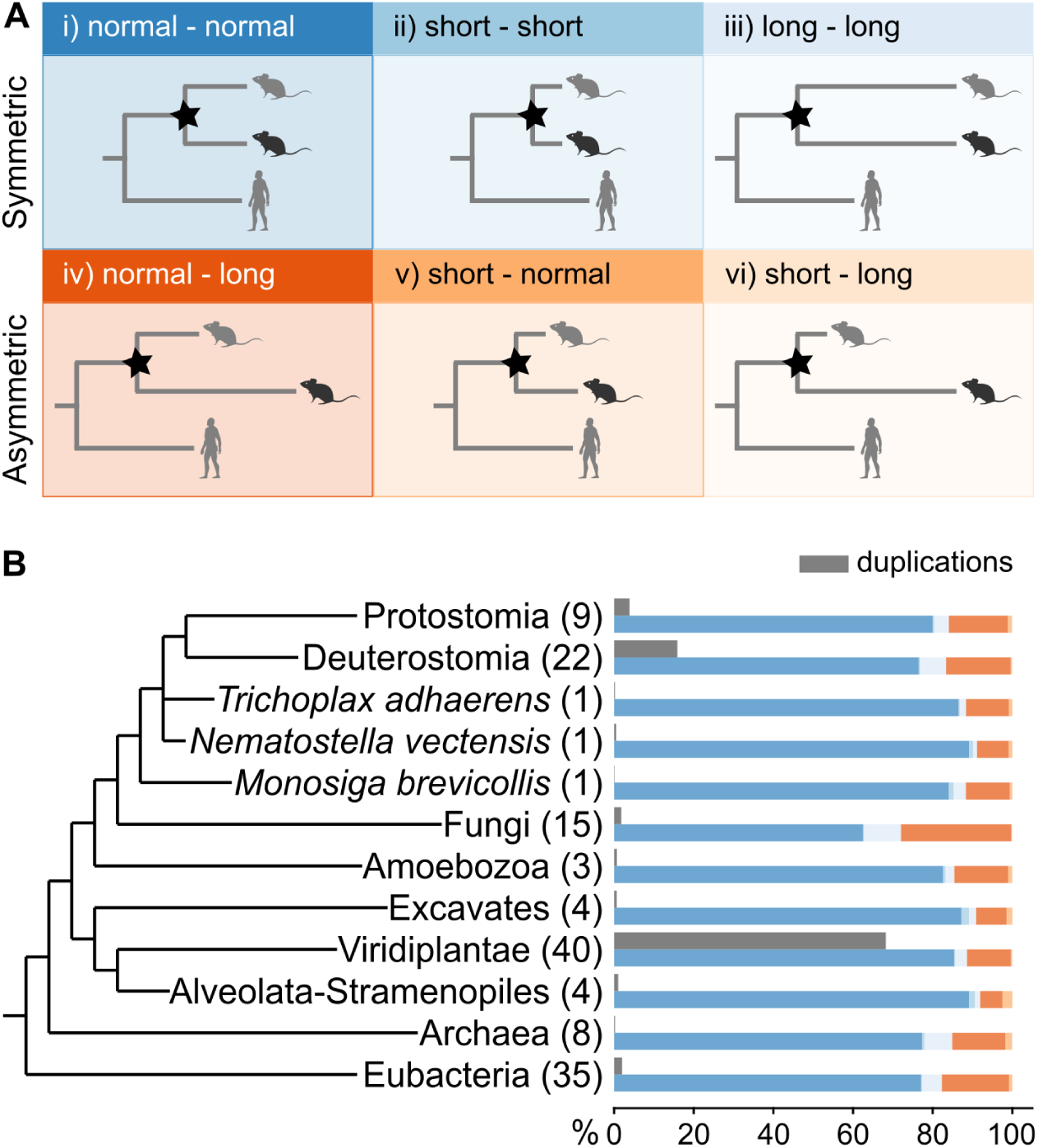
Branch length analysis following duplication events. **(A)** Depiction of the six categories indicating differences in branch length following a duplication event. The two rows divide the symmetric and asymmetric evolution of paralogues following duplication. The light silhouette (grey mouse) indicates the least diverged orthologue (LDO), and the dark silhouette (black mouse) indicates the most diverged orthologue (MDO). **(B)** Species tree showing the major lineages analysed. The numbers in brackets indicate the number of species included from the lineage. Grey bars represent the percentage of total duplications contributed by each lineage. Coloured bars indicate the proportion of duplications within each category (colour scheme as in Fig. 2A), relative to the total number of duplications in that lineage. The absolute numbers of duplications are provided in Supplemental Table S3.

The analysis gave a total of 926,814 duplication events across the Tree of Life, with an average of 68 duplications per family. When examining the major lineages (Fig. 2B, grey bars), the Viridiplantae lineage exhibits the highest percentage of duplications from the total number of duplications (68%, Fig. 2B, Supplemental Table S3), followed by Deuterostomia (16%) and Protostomia (4%). Internal nodes within this tree contributed only 5% of the total number of duplications (Supplemental Table S3). These findings highlight the significant role of gene duplication in the evolutionary history of plants, in agreement with previous studies (Jiao et al. 2011; Soltis et al. 2015).

Further analysis of the distribution of duplications across the six categories within each major lineage (Fig. 2B, coloured bars) revealed that the vast majority of duplications fell into the normal-normal group (∼82%). The second most abundant category was the normal-long group (∼13%), which includes paralogues with significantly different branch lengths. The remaining four categories (short-short, long-long, short-normal and short-long) accounted for only ∼6% of duplications. When grouping these categories based on their evolutionary pattern, 86% displayed symmetric evolution (both branches following similar rates: normal-normal, short-short and long-long), with 14% exhibiting asymmetric evolution (one significantly longer: normal-long, short-normal, short-long). These results indicate that following duplication events, the majority of paralogous pairs have similar evolutionary rates despite observed variations in absolute branch lengths.

Gene duplications in internal branches represent more ancient duplications compared to species-specific duplications (terminal branches). To investigate any differences, the distribution of duplications between these two types of duplications was examined within each major lineage subtree, excluding three major lineages which consist of only one species (*Trichoplax adhaerens*, *Nematostella vectensis*, *Monosiga brevicollis*). Across all cases, a large proportion of duplication events were present in terminal branches (∼70%, Supplemental Fig. S3A), which was similar in all major lineages (Supplemental Fig. S3B). When analysing the distribution of the duplications between the two evolutionary models, a larger proportion of duplications followed symmetric evolution (Supplemental Fig. S3C) in all major lineages (Supplemental Fig. S3D). Furthermore, the proportion of duplications following asymmetric evolution was higher in internal than in terminal branches, suggesting that ancestral duplications may have more time to evolve divergent gene copies.

### LDOs show greater conservation of protein structure

As protein structure is associated with protein function (Orengo et al. 1999), structural similarity can be used to test for functional divergence. However, structural similarity can be influenced by sequence similarity, as structural predictions often rely on sequence information. To investigate this relationship, structural similarity (measured by Foldseek LDDT) and sequence similarity (percentage of identity) were calculated between paralogues classified into symmetric and asymmetric categories. A stronger correlation between structural and sequence similarity was observed in symmetric duplications (PCC = 0.75), whereas asymmetric duplications showed a weaker correlation (PCC = 0.55) (Supplemental Fig. S4). This substantial difference between the two indicates that different evolutionary constraints or mechanisms act following symmetric versus asymmetric duplication.

To test the least diverged orthologue conjecture, controlling for sequence divergence, structural similarity was computed between each paralogous gene copy and a corresponding co-orthologous outgroup gene. The outgroups og1 and og2 were selected to have similar levels of sequence divergence to the MDO and the LDO, respectively (see Methods). To balance the need for comparable sequence divergence with the availability of suitable cases, the absolute difference in sequence identity between MDO-og1 and LDO-og2 was required to be at most 20%.

In asymmetric duplications, the median structural similarity for MDO-og1 pairs (0.723, dark orange) remained consistently lower than that for LDO-og2 pairs (0.7662, light orange) (Fig. 3A). In contrast, differences were minor for symmetric duplications (MDO-og1: 0.8506; LDO-og2: 0.8557). These findings suggest that although sequence similarity influences structural similarity, structural differences may persist beyond what sequence alone can explain, potentially reflecting functional divergence.

**Figure 3.**
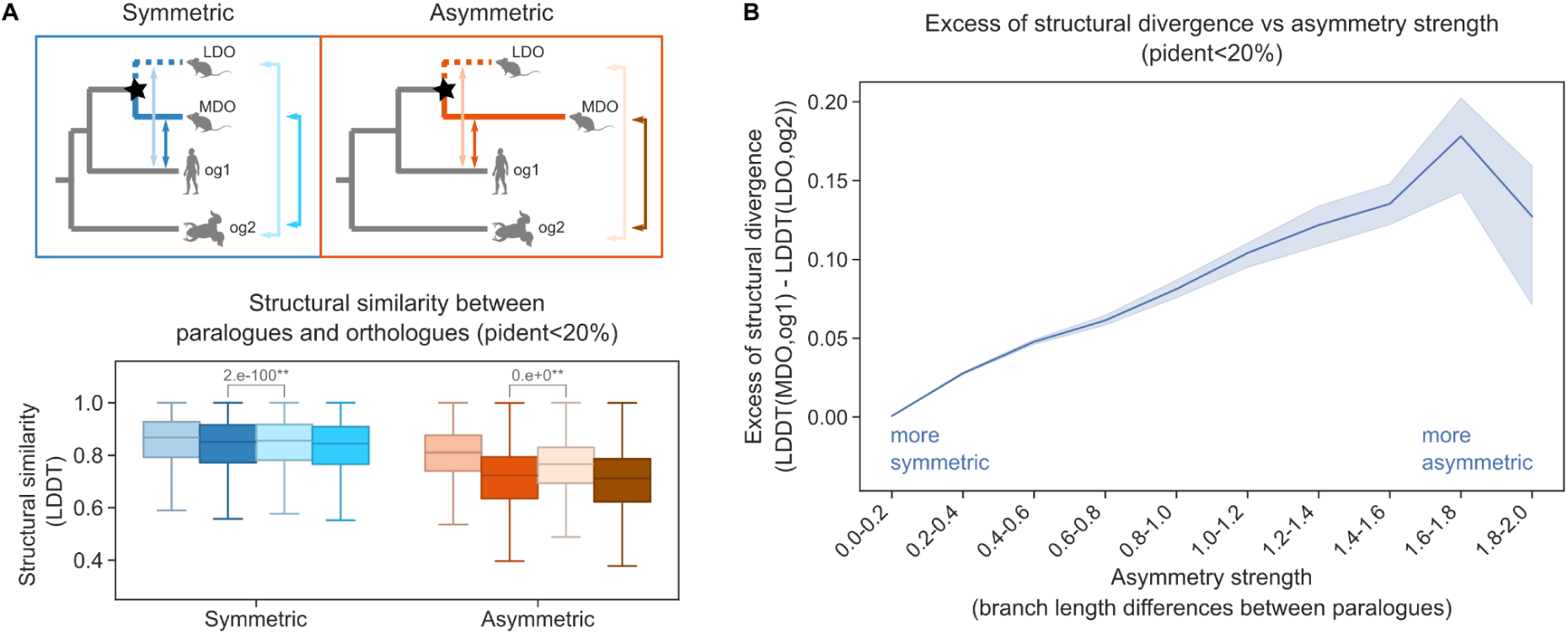
Structural similarity between paralogues and orthologues. **(A)** Structural similarity of each paralogue to two different orthologues (og1 and og2) (n=466,476). Box colours correspond to the arrows in the figure, indicating the comparisons made. The *p*-value is indicated on top of the figure for the Mann-Whitney *U* test (** *p*<0.01). All comparisons and *p*-values are indicated in Supplemental Table S4. **(B)** Comparison of the excess of structural divergence and the asymmetry strength (n=466,476). The x-axis represents increasing bins of asymmetry strength.

To further explore this, the asymmetry strength (measured as the observed branch length differences between paralogues) was compared to the excess structural divergence (calculated as LDDT(MDO, og1) − LDDT(LDO, og2)) (Fig. 3B). The analysis revealed that greater asymmetry was associated with a stronger structural divergence of the MDO copy. A positive correlation was observed up to an asymmetry strength of approximately 1.8 (Fig. 3B). The decline in the excess of structural divergence may reflect the retention of conserved structural motifs despite the large sequence divergence.

To ensure that these results were not influenced by the choice of sequence identity difference threshold, additional analyses were performed using different distance similarity thresholds between 5% and 50%. In all cases, similar trends were observed (Supplemental Fig. S5, S6). Taken together, these results indicate that increased asymmetry in sequence evolution between paralogues is associated with greater structural divergence from the outgroup, even after controlling for sequence similarity. This supports the use of protein structure as a proxy for functional divergence. In line with the LDO conjecture, asymmetric duplications exhibit greater functional divergence, with the LDO copy more likely to retain the ancestral function.

### Differences in branch length support differences in gene co-expression

To test functional differences across genes, gene co-expression was also used as a proxy for functional similarity. For animals, expression atlases of 16 species across 32 anatomical entities were obtained from the Bgee database (Bastian et al. 2021). For plants, publicly available expression atlases from 20 species across 72 anatomical samples were used (Julca et al. 2021; Koh et al. 2023) (Supplemental Table S5). In order to assess if the observed differences in branch length following gene duplication have functional implications, gene co-expression between paralogues resulting from symmetric and asymmetric evolution was compared using the Pearson correlation coefficient (PCC). Subsequently, the direction of change was investigated by comparing the expression profiles of each paralogue to their co-orthologous outgroup gene.

When analysing pairs of paralogues in each category, within both animals and plants, paralogues from the asymmetric category showed lower correlation values compared to those from the symmetric category (Fig. 4A). Furthermore, since most duplications were located in terminal branches and represent more recent duplication events, their gene expression profiles were evaluated separately (Supplemental Fig. S7). In both cases, terminal and internal branches present consistent results, with paralogues in the symmetric category showing slightly higher co-expression values. Notably, symmetrically evolving paralogues arising from duplications on the terminal branches within plants exhibited the highest co-expression (Supplemental Fig. S7). This observation may suggest that recent duplications have undergone less evolutionary divergence due to a shorter time frame. However, the same pattern is not observed in animals.

**Figure 4.**
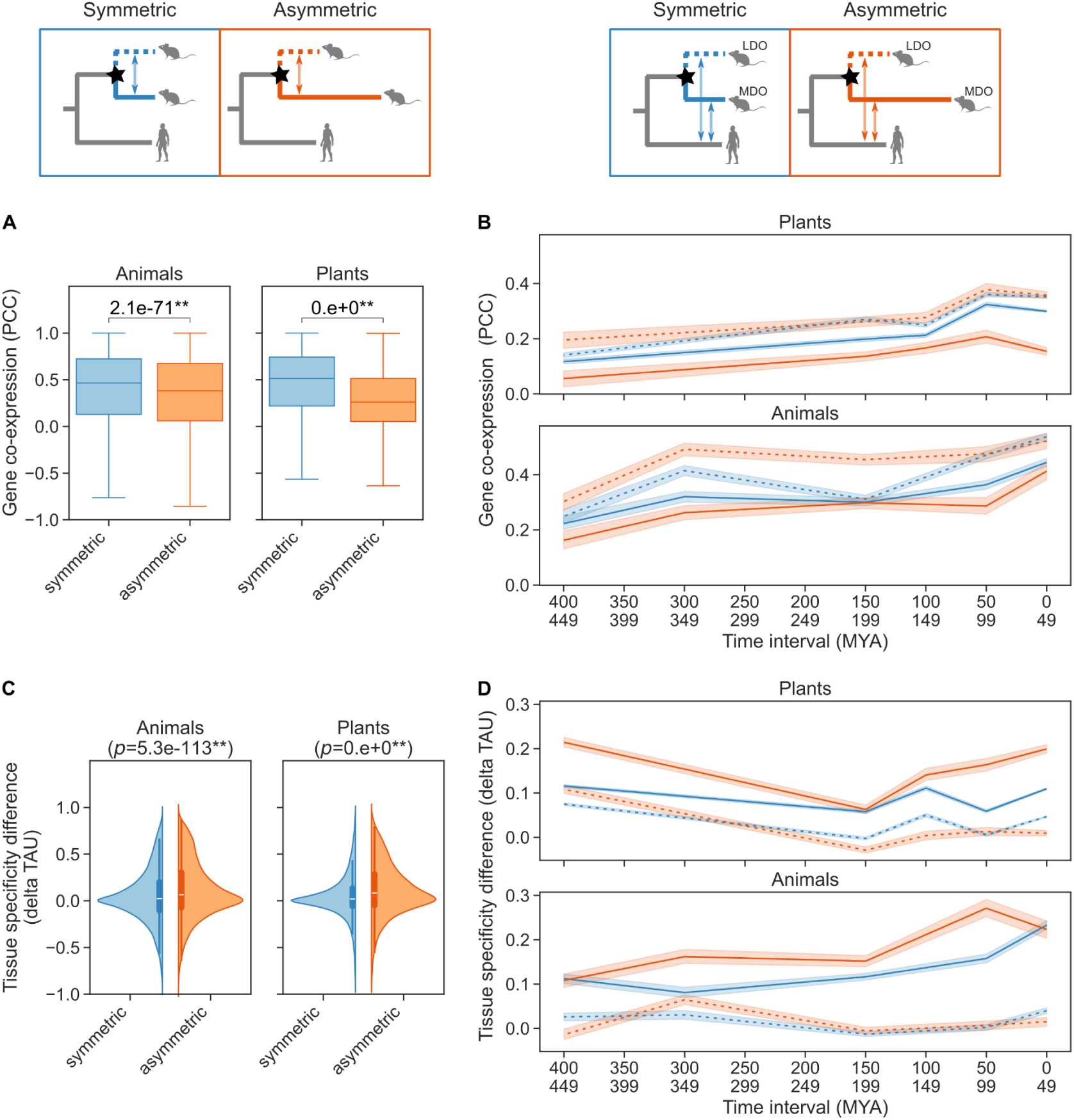
Functional analysis of paralogues using expression data. **(A)** Gene co-expression values of paralogues following symmetric (n=275,988) and asymmetric (n=31,671) duplications, calculated using the Pearson correlation coefficient (PCC). The *p*-values are indicated on top of the figures for the Mann-Whitney *U* test (* *p*<0.05 and ** *p*<0.01). **(B)** Gene co-expression (PCC) between the LDO (dashed lines) and MDO (solid lines) genes and their closest outgroup co-orthologue following symmetric (blue) and asymmetric (orange) duplications. The sample sizes for this plot are as follows: plants symmetric n=114,702, plants asymmetric n=13,533, animal symmetric n=17,493, and animal asymmetric n=7,490. **(C)** Violin plot showing the tissue specificity differences (delta Tau) between the paralogues following symmetric (animals n=77,889, plants n=275,988) and asymmetric (animals n=17,205, plants n=31,671) duplications. **(D)** Tissue specificity differences (delta Tau) between the LDO (dashed lines) and MDO (solid lines) genes and their closest orthologues following symmetric (blue) and asymmetric (orange) duplications. The sample sizes are the same as in (B). In plots (B) and (D), the x-axis indicates the divergence time between the species compared in time intervals. Plant and animal datasets were compared independently. Above the panels, the comparisons used are depicted.

The results may be influenced by the different species’ evolutionary history (for example, polyploid plants). Therefore, each species was analysed independently and showed consistent results (Supplemental Fig. S8). Amongst the animals, 13 species showed similar patterns, indicating lower co-expression values between paralogues within the asymmetric category. However, three species, *Felis catus*, *Drosophila melanogaster*, and *Gorilla gorilla gorilla*, did not show significant differences between the two categories. For plants, 19 species supported the previous pattern, whilst only one (*Amborella trichopoda*) did not. Moreover, the polyploid plants (*Brassica napus*, tetraploid; *Triticum aestivum*, hexaploid; *Glycine max*, tetraploid) exhibited major differences between symmetric and asymmetric paralogues. In summary, these results reveal a general trend where paralogues following asymmetric evolution exhibit less co-expression, suggesting that sequence divergence between paralogues correlates with differences in gene expression. Nevertheless, discrepancies observed in a few species highlight the importance of including many different species in this kind of study.

To investigate the direction of functional changes following duplication – whether the least (LDO) or most diverged orthologue (MDO) conserves the ancestral gene function – co-expression patterns were analysed between each paralogous gene copy and their co-orthologous outgroup gene. For plants, 112,015 duplications were analysed, with 99,995 symmetric and 12,020 asymmetric duplications. For animals, 22,885 duplications were examined, of which 16,273 were symmetric and 6,612 were asymmetric duplications. Then, within each category, gene co-expression was computed (PCC) between each copy and the outgroup gene (Fig. 4B). Differences were observed in both symmetric and asymmetric categories, with greater differences observed between the asymmetric category (Fig. 4B). Moreover, in both datasets – animals and plants – the gene following the longest branch (MDO in the asymmetric category), consistently displays low co-expression values (Fig. 4B), suggestive of a lower functional similarity with the ancestral gene. Similar results were observed by analysing each species independently (Supplemental Fig. S9).

In order to study the effect of time since duplication, co-expression was compared across species divergence time (Fig. 4B). In both animals and plants, more ancient paralogues exhibited a slight decrease in co-expression.

In summary, these results suggest that paralogous genes descending from duplications with significantly different branch lengths have more distinct co-expression values. In particular, the most diverged orthologues (MDO) display the greatest change in gene expression, thus suggesting that they have a greater functional divergence from the ancestral gene.

### Most divergent orthologues (MDO) show more functional specificity

Previous studies have shown that after gene duplication, paralogues tend to increase the levels of tissue specificity, which occurs promptly after the duplication event (Huminiecki and Wolfe 2004; Huerta-Cepas et al. 2011). To test whether the least diverged orthologue (LDO, paralogue with a shorter branch) or the most diverged orthologue (MDO, paralogue with a longer branch) has different tissue specificity, the Tau score (defined in Methods) was computed for each gene in every species.

To compare each category, the difference in tissue specificity was calculated (delta Tau) between the most diverged orthologue (MDO) and the least diverged orthologue (LDO) in each group (symmetric and asymmetric duplications, Fig. 4C). A value of zero indicates no difference in sample-specificity between the compared genes, while positive values indicate that the MDO exhibits more specificity and negative values signify that the LDO is more specific. The results of this analysis show that paralogues within the asymmetric category, in both animals and plants, have slightly higher positive values, indicating that the MDOs tend to have a more tissue-specific pattern of expression (Fig. 4C). Similar results were observed when analysing duplication events separately for internal and terminal branches, with terminal branches exhibiting more pronounced differences (Supplemental Fig. S10) and independently within each species (Supplemental Fig. S11). These results suggest that the gene copy which exhibits greater sequence divergence tends to have greater tissue specificity.

To determine whether the most diverged orthologue (MDO) also displays increased tissue specificity compared to the ancestral gene, the difference was computed between the Tau values for each paralogous gene copy and their co-orthologous outgroup gene (Fig. 4D). A zero value indicates no difference in tissue specificity, whilst positive values suggest increased specificity of the paralogous gene. In both categories, there are differences between the paralogues, with the MDO genes exhibiting higher delta Tau values. Specifically, in both plants and animals, the MDO gene of the asymmetric category consistently showed the greatest increase in tissue specificity over time (Fig. 4D). Moreover, similar results were observed when analysing individual species (Supplemental Fig. S12). In addition, the delta Tau in animals showed a slight decrease over time, whilst in plants this decrease was not apparent (Fig. 4D). This result indicates that tissue specificity in animals may decrease over time, with recent duplications exhibiting higher specificity. To summarise, these findings suggest that when one gene copy is under selective pressure to retain the ancestral function (LDO), the other with the longer branch (MDO) tends to acquire a more specialised role, which may differ over time.

### Well-characterised gene families corroborate the LDO conjecture

To explore the findings of this study, well-characterised gene families from the literature with available experimental data were examined. The analysis revealed that symmetric duplications (normal-normal category) generally retain similar functions between paralogues, whereas asymmetric duplications (normal-long category) are more likely to diverge functionally. In these asymmetric pairs, the least diverged orthologue (LDO) typically preserves the ancestral function.

A representative example is the *RNF113* gene family (Supplemental Fig. S13A), which duplicated independently in human and mouse. The human duplication is classified as normal-long, while the mouse duplication is normal-normal. In *Drosophila*, the orthologue *mdlc* is broadly expressed and essential for both neuronal development and spermatogenesis (Carney et al. 2013; Correia et al. 2024). In humans, the LDO gene *RNF113A* retains broad expression and the ancestral neuronal role (Tsampoula et al. 2022), while *RNF113B* (MDO gene) is testis-specific and essential for meiosis (Correia et al. 2024). In mouse, both *Rnf113a1* and *RNf113a2* retain broad expression, yet *Rnf113a2* (MDO gene) is required for spermatogenesis and *Rnf113a1* (LDO gene) for neuronal development (Correia et al. 2024), indicating some functional partitioning despite the classification as normal-normal.

Several clear examples of functional redundancy in normal-normal pairs were also identified. *SEC10a/b* in *Arabidopsis* (Supplemental Fig. S13B) share exocytosis-related functions (Vukašinović et al. 2014). The paralogues *Sar1a/b* in mouse (Supplemental Fig. S13C) are fully interchangeable for viability (Tang et al. 2024). For the paralogues *Cdx1/Cdx2* (Supplemental Fig. S13D), a knock-in of *Cdx2* rescues the *Cdx1*-null phenotype without defects (Savory et al. 2009).

In the normal-long category, *PRP18A/B* in *Arabidopsis* (Supplemental Fig. S13E) show functional divergence. *PRP18A* (LDO) retains a role in alternative splicing, similar to its yeast orthologue, whereas *PRP18B* (MDO) shows no detectable expression in two-week-old seedlings, and T-DNA insertion mutants exhibit no observable phenotype compared to wild-type plants (Kanno et al. 2018). The paralogues *LAZY1/LAZY6* in *Arabidopsis* (Supplemental Fig. S13F) also illustrate this pattern: *LAZY1* (LDO) controls shoot orientation as in rice, whereas *LAZY6* (MDO) has a different expression profile and no significant phenotype in mutants (Li et al. 2007; Yoshihara and Spalding 2017; Waite and Dardick 2024). Similarly, studies in knockout mice suggest that *Becn1* and *Becn2* (Supplemental Fig. S13G) have distinct functions in regulating autophagosome formation and mitophagy (Galluzzi and Kroemer 2013; Quiles et al. 2023).

Overall, these examples are consistent with the broader patterns described in this study. Nevertheless, certain cases, such as the *Rnf113* family in mice, highlight the complexity of functional evolution post-duplication and suggest that exceptions or intermediate scenarios exist. While general trends across the Tree of Life were the focus of this study, gene-specific dynamics should be considered in future analyses.

## Discussion

To investigate the least diverged orthologue (LDO) conjecture, a novel method was developed to identify paralogues with significantly divergent branch lengths, representing asymmetric evolution. This enabled the exploration of functional divergence between genes following duplication using gene trees from the PANTHER database, covering 143 species, along with structural data for more than one million proteins and expression atlases from 16 animal and 20 plant species. This study provides the most comprehensive analysis, using the largest dataset to date for investigating functional divergence following gene duplication events. Furthermore, it avoids the pitfalls associated with comparing Gene Ontology (GO) functional annotations across different species (Altenhoff et al. 2012; Thomas et al. 2012; Gaudet and Dessimoz 2017).

In the PANTHER database, pairs of extant genes are classified as least diverged orthologues (LDOs) based on their gene tree branch lengths immediately following each post-speciation duplication event (Mi et al. 2010). However, this definition uses only the observed branch length, with no test being employed to determine the significance of any differences between the branch lengths following duplication. To address this gap, this study provides a generalisation of the definition (least and most diverged orthologues – LDO and MDO), as well as a method to evaluate gene tree branch lengths and identify post-speciation duplication events with significant rate differences between branches. The results show that, despite observed differences in branch lengths between paralogues, most are not statistically significant (symmetric duplications). These results were consistent across all lineages included in this analysis. Previous studies have suggested that paralogues evolve at similar rates (Kondrashov et al. 2002) and approximately only 25% of duplications within a species display asymmetric evolution (Conant and Wagner 2003). Other studies have reported even lower occurrences, observing less than 20% exhibiting asymmetric evolution following duplication (Vance et al. 2022). A recent study observed that the most frequent fate of duplications is conservation (symmetry), followed by neofunctionalization (asymmetry) (Kalhor et al. 2024). The results presented here are consistent with previous studies, indicating that genes following a duplication event tend to evolve symmetrically. However, this observation does not preclude the possibility of functional changes occurring between paralogues.

If the least diverged orthologue conjecture holds true, it suggests that the least divergent copy (LDO) tends to retain the ancestral function, whilst the most divergent orthologue (MDO) is free to evolve a new function (Ohno 1970). Notably, in asymmetric duplications, the MDO copy exhibited more structural differences compared to the ancestral gene than the LDO. Even after accounting for sequence differences between paralogues and their orthologues, the MDO still showed an increase in structural divergence in asymmetric duplications. It has been observed that the function of a protein strongly depends on its structure (Orengo et al. 1999; Sotomayor-Vivas et al. 2022) and that an increase in structural similarity corresponds to an increase in functional similarity (Barrio-Hernandez et al. 2023). Additionally, while protein structure similarity can correlate with sequence similarity to some extent, it is well recognised that protein structures evolve more slowly than sequences. Moreover, structural changes may not always be proportional to sequence divergence due to compensatory mutations or structural constraints (Chothia and Lesk 1986; Gan et al. 2002; Illergå rd et al. 2009). Considering this, the results of this study suggest that sequence divergence, as reflected in gene tree branch lengths, correlates with functional divergence at structural level.

Genes that co-express are often members of similar biological processes and frequently share similar functions (Eisen et al. 1998; van Dam et al. 2018). In this context, co-expression analyses were used to test the LDO conjecture, showing that paralogous genes which have shorter branches (LDO) tend to co-express with the orthologous gene, with larger differences observed in genes following asymmetric evolution.

Previous studies which focused on the models of gene duplication retention have proposed that either one (Gu et al. 2005; Brunet et al. 2006; Panchin et al. 2010; Pegueroles et al. 2013; Pich I Roselló and Kondrashov 2014) or both copies (Huminiecki and Wolfe 2004; Scannell and Wolfe 2008) undergo a rapid period of accelerated evolution, followed by a return to pre-duplication levels. Therefore, it has been suggested that expression divergence between paralogues can occur rapidly after a duplication event (Gu et al. 2002; Guschanski et al. 2017). Additionally, it has been observed that the expression of duplicated genes tends to evolve asymmetrically, with one copy maintaining the ancestral expression profile (Gu et al. 2005; Panchin et al. 2010; Pegueroles et al. 2013) and that asymmetric sequence divergence correlates with asymmetric functional divergence (Blanc and Wolfe 2004; Zhang et al. 2004; Kim and Yi 2006). Moreover, the rate of protein evolution may be influenced by expression levels, with highly expressed genes evolving more slowly due to stronger evolutionary constraints, such as protein misfolding (Drummond et al. 2005; Drummond and Wilke 2008). Overall, this study strongly corroborates previous works, supporting a rapid change in expression profile following duplication. Furthermore, the results highlight that the copy with the shortest branch (LDO), which evolves more slowly under major constraints, tends to retain the ancestral gene expression profile, further supporting the least diverged orthologue conjecture.

Assuming that the LDO conjecture holds and the MDO copy has a more divergent function, one hypothesis proposes that the new function tends to be more specific (Li et al. 2005). To test this “specialisation hypothesis”, the tissue-specificity of each gene copy was analysed. This revealed that following either symmetric or asymmetric evolution, the MDO copy tends to show higher tissue specificity, particularly notable in the MDO of asymmetric duplications. Additionally, for animals, the levels of tissue-specificity in paralogues, when compared with their co-orthologues, appear to change over time, with the MDO tending to have a more specialised function in recent duplications. These results are consistent with previous studies, which showed that gene duplication leads to a rapid increase in tissue specificity (Huminiecki and Wolfe 2004; Huerta-Cepas et al. 2011) and that expression divergence between paralogues results in increased tissue specificity (Assis and Bachtrog 2015). Furthermore, young paralogues were reported to be highly tissue-specific and become more broadly expressed with divergence time (Kryuchkova-Mostacci and Robinson-Rechavi 2016).

Genomic positional information has also been proposed as an additional indicator of functional retention after gene duplication. Specifically, genes that retain their ancestral genomic positions (“positional orthologues”) may be more likely to maintain their original function (Dewey 2011). This raises the possibility that, following duplication, the least diverged orthologue (LDO) might not only be identified by sequence divergence but also by positional conservation. Although the present study did not incorporate positional information, considering both sequence and positional divergence could provide a more nuanced understanding of functional retention and represents an important direction for future work.

In conclusion, this comprehensive study using expression and structural aspects of function corroborates the least divergent orthologue (LDO) conjecture: the gene copy with the shortest branch after a duplication event tends to retain the ancestral function. The LDO gene maintains a similar structure, expression profile, and tissue specificity to the ancestral gene, highlighting its functional conservation over time. Furthermore, the differences in structure and gene expression observed in paralogues following symmetric evolution suggested a rapid change occurring after gene duplication. Overall, these results significantly contribute to the understanding of gene duplication dynamics and their effect on gene function.

## Methods

### Gene tree processing

To study the least diverged orthologue conjecture, the species tree for 143 organisms and 15,693 gene trees were downloaded from PANTHER v18.0 (Thomas et al. 2022). PANTHER gene trees are annotated with evolutionary information in their internal nodes, including speciation, gene duplication, horizontal gene transfer (HGT), or unknown events (nodes for which the type cannot be reliably inferred). Specifically in bacteria, HGT events appear to be underestimated in this dataset due to the methodology and the type of data used (single reference genome per bacterial species rather than a pangenome). Some specific branches in the gene trees were excluded from the analysis. These include branches with poorly fitting distances (arbitrarily set to 2.0 substitutions/site in PANTHER), and some duplication events that involve: duplications at the root of the gene tree (no speciation event to compare); those with horizontal gene transfer or “unknown” events annotated on either side of the duplication; as well as those with gene losses immediately following the duplication. Finally, a total of 15,577 gene trees were used for further analysis (Supplemental Table S2).

### Calculation of the gene tree expected branch lengths

Expected gene tree branch lengths were calculated for 2,648,252 branches across 15,577 gene trees using a simple evolutionary model: the expected length between two speciation events in a gene tree is equal to the corresponding branch in the species tree scaled by a gene family-specific rate. In the case of gene loss, the model assumes that the branch length in the gene tree represents the sum of all the branches corresponding to the branch in the species tree scaled by the family-specific rate (Fig. 1A).

By expressing this as a system of approximate linear equations, it is possible to optimise the family rates and species tree branch lengths, constrained by the observed branch lengths within the many gene families. That is, 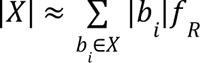, where *X* represents a specific branch in a gene tree (|*X*| being the length of that branch), which corresponds to a set of branches in the species tree, each denoted *b*_*i*_ (with length |*b*|), and *f* representing the family-specific factor.

The species tree used in PANTHER only provides topological relationships. As such, both the species tree branch lengths and family-specific factors were estimated from the observations of gene tree branch lengths. The method consists of two main steps:

1. *Species Tree Branch Fitting:* An iterative approach was used to fit the branch lengths of the species tree from the system of equations, whilst also optimising the family-specific factors. Initially, species tree branches were set to the median of the lengths of matching branches from all the gene trees. This way, the species tree branch lengths reflect the typical amount of genetic change observed across all genes. Using the initial species tree branch lengths, as described above, the family-specific factors were initialised according to the equations. The iterative procedure then has two steps: (1) refining the species tree branch lengths, using a least-squares solution, by estimating them whilst fixing the family-specific factor; (2) calculating the family specific factors, whilst keeping the species tree branch lengths fixed. Whilst optimising the species tree branches, an additional normalisation constraint was included, so that the average family-specific factor is 1. Convergence was achieved when the cosine distance between consecutive estimates is less than ^−9^. The final estimate of species tree branch lengths was rounded to 3 decimal places. The species tree, including estimated branch lengths, is available in both Newick and PNG formats in the results folder of the git repository on GitHub (https://github.com/DessimozLab/ldo_study/tree/main/results) and the Supplemental Scripts.
2. *Family-Specific Factors*: Using the final estimate of species tree branch lengths, family-specific rates were calculated, without the additional normalisation constraint used in step (1). The resulting family-specific factors are available in Supplemental Table S1.

In this way, the family-specific factor represents the best-fitting scaling factor that aligns the branch lengths of a gene tree with those of the species tree. It captures the overall rate of molecular evolution for each gene family, reflecting both neutral substitutions at tolerant sites and the effects of purifying selection at constrained sites. A family-specific factor greater than one indicates that the gene family is evolving faster — suggesting fewer functional constraints — while a factor less than one indicates slower evolution, consistent with stronger functional constraints compared to the average across families.

Using these species tree branch lengths and family-specific factors, expected branch lengths were calculated in the gene trees, before identifying statistically significant rate shifts (Supplemental Table S2). The Python scripts generated for this method are available on GitHub (https://github.com/DessimozLab/ldo_study) and in the Supplemental Scripts.

### Identification of branch length differences after duplication

Since all paralogues resulting from a duplication event can be classified as either least diverged orthologues (LDO, shorter branch) or most diverged orthologues (MDO, longer branch), a statistical method was developed to determine whether the differences observed in the branch length post-duplication were significant.

For 15,577 gene trees, the difference between observed and expected branch lengths (calculated under the model described in the previous section, from speciation event to speciation event) was computed (Fig. 1A). Then, to assess the statistical significance of the observed excess or depletion, these differences were converted into standardised *Z*-scores using the zscore function from the SciPy Python package v1.13.0 (Virtanen et al. 2020). An excess substitution can be interpreted as a potential signal of branch-specific adaptive selection and/or relaxed constraints. The resulting distributions of the normalised differences between branches with (876,261) and without (1,771,991) duplication were compared using the Mann-Whitney *U* test (also using SciPy). All further analysis filtered out gene trees without duplication events, which resulted in 13,597 gene trees and 364,829 duplication events distributed across different lineages of the Tree of Life.

For each duplication event, the two branches (Figure 1A) were compared using their *Z*-score of the difference between observed and expected (from speciation to speciation event) by computing *p*-values and using a two-tailed test (with significance level α=0.05) to identify significantly shorter and longer than expected branches. When more than two gene copies were present, indicating either successive duplications, multifurcation (that is, duplication nodes with more than two descendant branches), or both, the analysis was performed at every node starting from the leaves (Supplemental Fig. S14). Then, the least diverged orthologue (LDO) was identified and compared to the other paralogues. This approach ensures that a long branch is included only once in the analysis.

Based on these comparisons, duplication events were placed into six categories (Fig. 2A): i) normal-normal: both branches are not significantly different; ii) short-short: both branches are significantly shorter than expected; iii) long-long: both branches are significantly longer than expected; iv) normal-long: only one branch is significantly longer than expected; v) short-normal: only one branch is significantly shorter than expected; vi) short-long: one branch is significantly shorter than expected and the other is significantly longer than expected. Using this information, the dataset was divided into two duplication models: symmetric (normal-normal, short-short, long-long) and asymmetric evolution (normal-long, short-normal, short-long). All subsequent analyses were based on this classification into these two categories: symmetric and asymmetric duplication events.

### Pairwise comparisons of structural and sequence identity

Structural data for 1,829,120 proteins out of 1,968,858 of the PANTHER dataset were downloaded from the AlphaFold Protein Structure Database v4 (Varadi et al. 2022). The 139,738 remaining proteins did not have structural predictions available. Corresponding amino acid sequences for all proteins with available structural data were obtained from the UniProt database using the REST API (downloaded on 19 March 2025).

Pairwise structural similarity was computed for 926,814 paralogue pairs using Foldseek version 8.ef4e960 (van Kempen et al. 2024). The Foldseek LDDT (Local Distance Difference Test) score reports the average LDDT of the alignment. LDDT scores express the percentage of inter-atomic distances and range from 0 (not conserved distances) to 1 (perfect model) (Mariani et al. 2013).

To estimate sequence similarity, multiple sequence alignments were generated for each gene family using MAFFT v7.526 (Katoh and Standley 2013). From these alignments, pairwise percentage identity between paralogues was calculated. The Pearson correlation coefficient (PCC) was then computed to assess the relationship between sequence and structural similarity, separately for symmetric and asymmetric duplication categories.

In addition, pairwise sequence and structural similarity were calculated for each paralogue and their co-orthologous outgroup gene (857,663 comparisons), using the approach described above. Two outgroup genes were selected to account for differences in the absolute branch length of each paralogue. For the first outgroup (og1), co-orthologous genes were selected from the closest outgroup species before the duplication event in the gene tree. Then, the closest gene in that species (least diverged copy) was selected.

For the selection of the second outgroup (og2), the aim was to minimise the differences between LDO-og2 and MDO-og1, such that the sequence differences between MDO and og1 would be comparable to the differences between LDO and og2. The branch distance between MDO and og1 was calculated, and then an external orthologue with a similar distance to the LDO copy was selected as og2. To also account for sequence divergence, pairwise percentage identities between MDO and og1 and LDO and og2 were calculated using the multiple sequence alignment generated before (Supplemental Table S6).

For the analysis using the second outgroup (og2), the ideal scenario is that the difference in sequence similarity between MDO-og1 and LDO-og2 would be zero. In such cases, the comparison between MDO-og1 and LDO-og2 would be perfectly balanced in terms of sequence or branch distance. However, applying this strict filter drastically reduced the dataset size (from 548,818 to 1,319 (0.2%)). To address this, six more relaxed thresholds were used to filter for differences in percentage identity of: <5%, <10%, <20%, <30%, <40%, and <50%. Distributions of the different datasets were compared using the Mann-Whitney *U* test.

### Asymmetry strength and excess of structural divergence analysis

To test whether the observed branch length differences between paralogues (referred to as asymmetry strength) correlate with structural divergence, the branch length differences for both symmetric and asymmetric duplications were calculated. Then, the differences in structural similarity (LDDT) between the comparisons LDO-og2 and MDO-og1 were computed (Supplemental Table S7). This analysis was performed for all datasets generated using the different sequence similarity and branch length thresholds described in the previous section.

### Compilation of gene expression atlases

As another functional characteristic of paralogous genes, expression atlases of 16 animals and 20 plants were used. For animals, RNA-seq gene expression data and anatomical annotations (UBERON terms) were retrieved from Bgee 15.1 (Bastian et al. 2021). This consisted of 2,639 RNA-seq experiments across 32 different anatomical samples. For plants, expression atlases were taken from previous studies (The Plant Expression Omnibus) (Julca et al. 2021; Koh et al. 2023), and anatomical entities were obtained from the sample names of RNA-seq experiments in the NCBI Biosample database. Manual curation was conducted to define anatomical entity names (for example, leaf, flowers), resulting in 72 different anatomical samples for plants. All species included, for both animals and plants, had at least four different anatomical samples (Supplemental Table S5). Also, only genes with TPM >=2 in at least one sample were retained (plants = 446,852 and animals = 167,106 genes). Then, the TPM data of each gene was log-transformed using the hyperbolic arcsine function (Johnson and Krishnan 2022): *arcsinh(x)* = *ln*(*x* + √*x*^2^ + 1. Finally, the average value was computed for samples coming from the same anatomical entity.

### Gene co-expression analysis

To test for functional differences between pairs of paralogues (animals 95,094, and plants 307,659 pairs), as well as each paralogue and the closest orthologous outgroup (animals 24,983, and plants 128,235 pairs), gene-gene co-expression patterns were computed using the Pearson correlation coefficient (PCC). To facilitate comparison across different species, only equivalent anatomical entities were used (for example, leaf for *Arabidopsis* and tomato) (Supplemental Table S5). When selecting an orthologous outgroup to compare the paralogous genes to, the closest species was used for which data was available (Supplemental Table S8). Additionally, species were removed from the outgroup analysis when there were too few orthologous genes (<50 genes). Finally, distributions of the different datasets were compared using the Mann-Whitney *U* test.

### Analysis of gene expression specificity

Sample-specificity of genes, based on expression data, was calculated using the Tau score (Yanai et al. 2005; Kryuchkova-Mostacci and Robinson-Rechavi 2017). Tau values range from 0 to 1, where 0 indicates a gene is broadly expressed and 1 denotes tissue-specific expression. Gene pairwise comparisons were performed by calculating the difference in tissue specificity (delta Tau) between the two genes. Comparisons were performed for the pairs of extant paralogous genes (animals 95,094, and plants 307,659 pairs), as well as independently for each gene copy with the chosen orthologous outgroup gene (animals 24,983, and plants 128,235 pairs). Differences in their distribution were assessed using the Mann-Whitney *U* test.

### Estimation of divergence times

Divergence times (adjusted times) between species were obtained from the TimeTree database (Kumar et al. 2022) and are expressed in millions of years ago (MYA). These time estimates were used to plot the differences between genes in the expression profile and sample specificity analyses.

## Acknowledgements

This work was funded by the SNSF grant 205085. I.J. acknowledges support by a Young Investigator Grant from the Novartis Foundation for Medical-Biological Research (24C173). We thank all members of the Comparative Genomics lab and Riccardo Delli Ponti for their valuable insights. We also thank the reviewers for their positive feedback. Author Contributions: CD and PDT conceived the project idea. CD, AWV, NG, and IJ designed the methodology. AWV developed the scripts to analyse the data. AWV and IJ analysed the data. IJ and AWV wrote the manuscript with comments from all authors. All authors read and approved the final manuscript.

